# “Guess Who?”: Forensic genetics and archaeology converge to identify Cristopher Columbus descendants

**DOI:** 10.64898/2025.12.16.694569

**Authors:** I Navarro-Vera, J Yravedra Sainz de los Terreros, A Bonilla, M Tirapu, M Albert, P Jímenez, D Herránz, C García

## Abstract

The generation of DNA profiles from skeletal remains is a core challenge in forensic genetics, particularly for the identification of missing persons and the resolution of cold cases. Factors such as diagenesis, DNA fragmentation, and the presence of PCR inhibitors severely compromise traditional methodologies, limiting the potential for successful identification.

Similarly, in many archaeological studies, the presence of multiple individuals within a single burial site presents a significant challenge for accurate identification. This study shows how the most innovative forensic genetic techniques can be integrated into archaeological research, enabling the identification of individuals through the establishment of familial relationships between remains recovered from an archaeological site. A large-scale SNP assay, Forenseq® Kintelligence HT, has been applied to a familial group dating back to the 16th-17th century, allowing robust kinship calculations and lineage studies, that, in a first stage, lead to the identification of four individuals spanning four generations within the same family.

## INTRODUCTION

The archaeological research of a family pantheon founded in the 16th century, containing several buried members of Cristopher Columbus descendants, provided a unique opportunity for this study. The preset work outlines the second phase of the comprehensive investigation underway on the remains recovered from this family burial, located inside the church of Santa María de Gracia in the municipality of Gelves (Sevila, Spain) [1]. Access to the crypt has been guarded by ecclesiastical authority and was only recently permitted for the present archaeological investigation. Upon obtaining the necessary permits, the research team was able to conduct excavations and collect bone samples from various individuals within the crypt.

The present work focusses specifically on four individuals designed as I4, I13, I21 and I22. All these individuals dated to the 17th-18th century, pointing out to a specific branch of the family, the Counts of Gelves (direct descendants of Christopher Columbus), reflected in the historical records. Skeletal samples were collected for genetic analysis to conduct kinship studies, aiming to establish relationships to achieve accurate identifications.

The study of unidentified human remains (UHR) is one of the most critical functions of forensic science, [2,3]. When biological evidence is limited to skeletal remains, the identification process faces important molecular challenges. The main obstacle is diagenesis, the set of post-mortem processes that alter bone composition and degrade endogenous DNA. This results in severe DNA fragmentation, in which long nucleotide chains are broken into tiny fragments that are often too short to be analyzed using conventional forensic methods [3]. Additionally, skeletal remains often contain high concentrations of PCR inhibitors co-extracted from the environment, which interfere with DNA amplification [4]. These limitations mean that traditional methods based on short tandem repeat (STR) analysis, requiring relatively long DNA fragments, frequently fail to generate a complete, usable genetic profile.

In this context, the advent of Massively Parallel Sequencing (MPS), has represented a paradigm shift [5-9]. By targeting DNA segments shorter than 150 base pairs, MPS-based assays dramatically increase the likelihood of obtaining informative genetic profiles even from highly degraded bone samples.

The accumulated evidence conclusively demonstrates the superior performance of Massively Parallel Sequencing (MPS) kits with short amplicons targeting SNP markers, such as the ForenSeq® Kintelligence kit (QIAGEN), in degraded samples. SNPs are preferable to STRs for the analysis of skeletal remains for one fundamental reason: the required amplicon size. Whereas traditional STR kits may require DNA fragments of up to 400–500 bp, modern SNP panels such as ForenSeq® Kintelligence are designed with drastically shorter amplicons, in this specific case with 97.8% of them below 150 bp [4,5]. This feature exponentially increases the probability of successful amplification of a marker from the fragmented DNA that typifies bone.

This study demonstrates how the application of a high-throughput sequencing technique based on SNP genotyping, such as Kintelligence (encompassing ∼10K SNPs), has enabled the acquisition of highly valuable genetic profiles out of skeletal remains proceeding from four individuals that date back to the 17th and 18th-century.

The genetic data obtained allowed the establishment of kinships among the four individuals founded in the crypt, with likelihood ratio (LR) values providing sufficient evidence to build a consistent family pedigree leading to an accurate identification. Furthermore, the analysis of SNPs on the sex chromosomes has strengthened these conclusions, enabling the reconstruction of a paternal lineage spanning four generations.

## MATERIAL AND METHODS

### Skeletal samples

The skeletal remains of all individuals were commingled and distributed across four unidentified boxes, that were labelled as Box 1-4. Following the anthropological study, the minimum number of individuals (MNI) was established at 12, with biological sex estimation indicating 6 males and 6 females. I4 was located alone in one Box 3, I13 was in Box 1, which held remains corresponding to eight different individuals according to the MNI determined by the anthropological study. Box 4 contained three individuals, including I21 and I22. Individuals were classified based on age estimation, stature, bone pathologies, and information derived from historical records.

Dental and molar specimens were retrieved from the individuals included in this study. Efforts were made to select those with optimal preservation. Specimens that were still anchored to the mandibular arch were chosen, ensuring they showed no visible fractures or signs of previous infections.

### DNA extraction and quantification

After cleaning and disinfection, bone samples were grinded using a Tissuelyser II (QIAGEN). Bone powder was incubated O/N in a lysis/decalcification buffer following previously described methods [10]. Subsequent DNA extraction was carried out using the large volume protocol in an EZ2 Connect instrument (QIAGEN). DNA concentration process was carried out using Amicon®Ultra-0.5 columns (Millipore). The obtained DNA was quantified by real-time PCR using Quantifiler™Trio DNA Quantification kit in a QuantStudio5 thermocycler (ThermoFischer).

### Library preparation and targeted sequencing

500 pg DNA input of each sample were used for library preparation following the manufacturer’s protocol of the ForenSeq® Kintellingence HT kit (QIAGEN). DNAse/RNAse free water was used as negative amplification control and 1ng of the genomic DNA control sample NA24385 (HG002-Coriell Institute for Medical Research, Camden NJ) provided with the kit, was included as positive control. Both were processed together with the test samples. To assess the size, quantity, and integrity, of the obtained libraries, these were quantified using both a fluorometric (Qubit™) and an electrophoretic method (TapeStation™). Subsequently, libraries were diluted to 0.45 ng/µL each and pooled together. A total of 8 libraries were pooled together: 6 sample libraries, positive and negative control. The denaturalized pool was loaded on a MiSeq® FGx Reagent Cartridge. MPS run was set up in a MiSeq®FGx instrument. Targeted sequencing took place on a Standard MiSeq®FGx Flow Cell (QIAGEN).

### Data analysis using the UAS

Sequenced libraries were visualised and analysed using the ForenSeq® Kintelligence Analysis Module in the ForenSeq® Universal Analysis Software (UAS) v2.7(QIAGEN). Allele calling was performed as described in the ForenSeq® Kintelligence Kit Reference Guide and the Universal Analysis Software – Kintelligence HT Module Reference Guide [11,12]. All analysis within the UAS were performed with the default analytical and interpretation thresholds at 1.5 % in the ForenSeq® Kintelligence HT analysis method. This represents a minimum read count of 10 reads [13]. Kinship calculations were carried out with the UAS associated local database, applying the manufacturer’s default settings.

## RESULTS

DNA proceeding from combined extractions for each sample was quantified. Highly satisfactory results were obtained, showing a good preservation of the genetic material along the years and during the extraction process. Enough DNA was gained in order to perform the subsequent MPS assay with the input recommended by the manufacturer. Degradation index was lower than expected among the samples, with exception of the one corresponding to I22, that showed a significantly higher DI value (table 1).

**Table 1:** Quantification data. Target amplicon sizes. Large autosomal 214 bp, Small autosomal 80 bp and Y-chromosome 75 bp.

| | DNA quantification (ng/μL) | | | |
| --- | --- | --- | --- | --- |
|  | I13 | I21 | I4 | I22 |
| Large autosomal | 0,018 | 0,132 | 0,075 | 0,001 |
| Small autosomal | 0,153 | 0,140 | 0,081 | 0,025 |
| Y chromosome | 0,145 | 0 | 0,097 | 0,012 |
| Degradation Index | 8,5 | 1,1 | 1,1 | 43,6 |

In terms of sequencing, run metrics such as cluster density, passing filter, phasing and prephasing values were in the expected range (1018k/mm^2^, 91.23%, 0.159% and 0.045% respectively). Positive control showed 2.9·10^6^ reads and concordance with the expected genotypes. Both, library preparation negative control and DNA extraction negative control resulted in no reads. The Human Sequencing Control (HSC) used as internal control by the UAS was in the expected quality range and intensity. Samples also showed a good performance, achieving very good parameters in terms of number of reads as well as the number of SNPs typed (table 2). A direct correlation between DNA quantification/DI values and the number of reads and SNPs typed has been observed, with sample I22 yielding the lowest results.

**Table 2:** Kintelligence HT results. Total reads: number of reads per sample; Total SNPs: number of SNPs typed: X/Y-SNPs-SNPs located in X/Y Chromosome; aSNPs ancestry SNPs; phSNPS phenotype SNPs; iSNPs: identity SNPS and kSNPs: kinship SNPs Typed markers are expressed as total number (Nr) and percentage of markers (%) typed.

| Sample ID | Total reads | Total SNPs |  | X-SNPs |  | Y-SNPs |  | aSNPs |  | phSNPs |  | iSNPs |  | kSNPs |  |
| --- | --- | --- | --- | --- | --- | --- | --- | --- | --- | --- | --- | --- | --- | --- | --- |
|  | Nr | Nr | % | Nr | % | Nr | % | Nr | % | Nr | % | Nr | % | Nr | % |
| Ind. 4 | 3.109.216 | 10123 | 99 | 104 | 98,1 | 84 | 98,8 | 56 | 100 | 24 | 100 | 93 | 98,9 | 9764 | 99 |
| Ind. 13 | 1.004.854 | 9339 | 91,3 | 89 | 84 | 67 | 78,8 | 48 | 85,7 | 24 | 100 | 82 | 87,2 | 9031 | 91,5 |
| Ind. 21 | 3.004.767 | 10033 | 98,9 | 106 | 100 | 0 | 0 | 55 | 98,2 | 24 | 100 | 93 | 98,9 | 9757 | 98,9 |
| Ind. 22 | 338.513 | 6814 | 66,6 | 63 | 59,4 | 37 | 43,5 | 36 | 64,29 | 10 | 41,7 | 55 | 58,5 | 6615 | 67,1 |

The observed read depth was variable between samples and alleles. A correlation between the number of alleles typed together with a higher read depth and the quality of the extracted DNA of the samples was observed. Samples I4 and I21 showed the highest number of highly covered alleles. In both cases, over 80% of the typed alleles had a coverage over 50x, meaning 12169 and 11983 alleles respectively; only a 3% and a 4%, showed the lowest coverage ranging from 10-20x. Samples I22 and especially I13 showed lower allele coverage values, yet still typing over 70% and 88% of the SNPs at mor than 20x (figure 1). Nevertheless, allele coverage values were considered very satisfactory overall, since an average of 88% of the markers showed coverage values over 20x. These metrics allowed to get confident genotypes and proceed to kinship calculations with all the samples.

**Figure 1:**
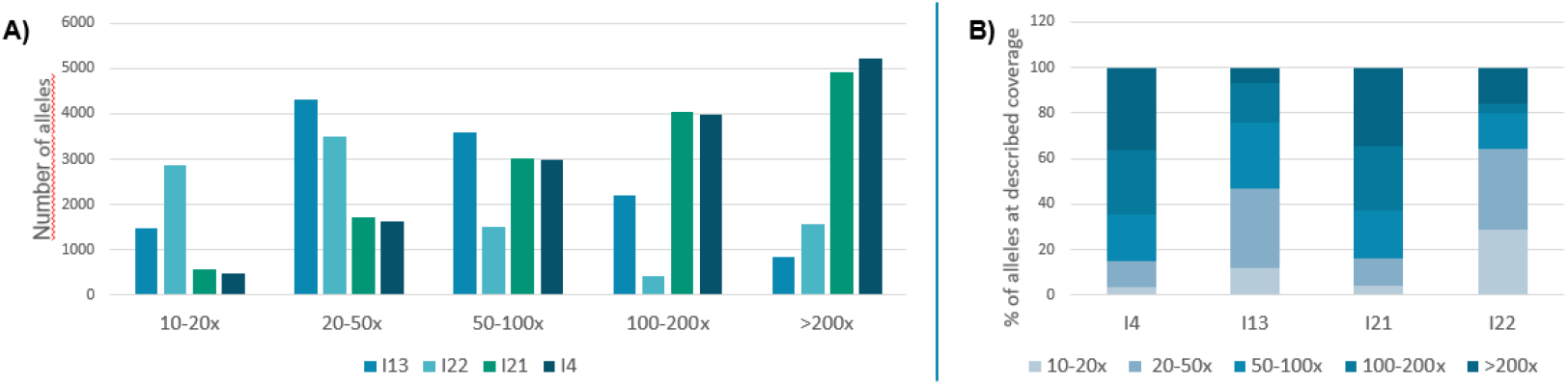
Typed alleles and coverage. **A)** Number of alleles typed clasified according to the coverage in the four different individuals. **B)** Perecentage of alleles typed at the different coverage ranges per sample.

After reviewing the overall metrics and the sample results, kinship analysis was carried out using the local database algorithm in the UAS v2.7 (QIAGEN). This algorithm performs pairwise calculations and is sensitive to the SNP overlap, meaning the number of typed SNPs in common between 2 samples, when searching for relationships. A Kinship coefficient indicating the degree of relatedness between two samples is computed if a match is found.

As expected in a familial group, all the individuals studied, showed different degree relationships to each other (table 3). But, in this particular case, some of the estimated relationship degrees were higher than initially expected.

**Table 3:** Kinship degree between the different individuals. Kinship coefficient calculated by the UAS algorithm is shown in blue scale. Kinship degree associated to the coefficient is shown in green scale.

| Individuals | I4 | I13 | I21 | I22 |
| --- | --- | --- | --- | --- |
| I4 | self | 0,067 | 0,119 | 0,178 |
| I13 | 3rd | self | 0,247 | 0,257 |
| I21 | 3rd | 1st | self | 0,253 |
| I22 | 2nd | 1st | 1st | self |

Besides the kinship calculations, SNPs located in the sexual chromosomes were analysed and compared. Firstly, all Y-Chromosome SNPs typed for all the male individuals (I4, I21 and I13) were coincident, pointing out to a common patrilineal ancestry. I4 and I13 shared the same genotypes in all the overlapping Y-chromosome SNPs (67) and both had coincidence with the 37 typed markers from I22 (data not shown).On the other hand, X-Chromosome SNPs were compared between the samples in order to rule out the compatibility as mother (1^st^ degree relationship) between sample I21 (the only female) with I22 and I13, given that, both of them pointed to a first degree relationship to her. After comparison, several inconsistencies were detected between I21 and I13, which disputes the firstly obtained kinship results between them. On the contrary, the overlapping X-chromosome SNPs between I21 and I 22 (63) were compatible with a first-degree relationship (data not shown).

Considering the data presented so far on kinship calculation based on autosomal markers, along with studies of SNPs located on both the Y chromosome and the X chromosome, and taking into account the historical documentation and anthropological data gathered on the individuals studied, the possible kinship hypotheses were formulated. The UAS algorithm was used to calculate logLR values based on the proposed kinship hypotheses as well as the total amount of shared DNA between each pair of related individuals. The longest stretch of shared DNA between each related pair was also estimated (table 4). The obtained values in terms of logLR and shared DNA were highly consistent and enabled the establishment of a clear family genealogical tree (figure 2).

**Table 4:** Kinship study results. Shows the mam results obtained out of the UAS local database algorithm. SNP overlap: number of SNPs in common between two samples. Shared DNA Total amount of shared DNA between two samples çM Centimorgans, Longest shared segment The longest street) of shared DNA between two samples. UAS relationship degree: relationship degree established by the UAS algorithm. Established relationship degree relationship degree established based on documentation and genetical findings in this sudy. Relationship H1. kinship hypothesis used, for logLR calculations. logLR. base-10 logarithm of the ILikelihood Ratio, based on the given kinship.

| Sample ID | Related Sample ID | SNP overlap | shared DNA (cM) | Longest shared segment (cM) | UAS relationship degree | Established relationship degree | Relationship H1 | log10 LR |
| --- | --- | --- | --- | --- | --- | --- | --- | --- |
| I4 | I13 | 9076 | 570,257 | 121,503 | 3rd | 3rd | Great-Grandfather | 42,63 |
|  | I21 | 9771 | 1498,744 | 160,103 | 3rd | UNK | N/A | N/A |
|  | I22 | 6648 | 1196,556 | 194,555 | 2nd | 2nd | Grandfather | 151,36 |
| I13 | I4 | 9076 | 570,257 | 121,503 | 3rd | 3rd | Great-Grandson | 42,63 |
|  | I21 | 9077 | 2185,016 | 194,391 | 1st | 2nd | Grandson | 361,99 |
|  | I22 | 6538 | 384,682 | 134,761 | 1st | 1st | Son | 410,64 |
| I21 | I4 | 9771 | 1498,744 | 160,103 | 3rd | UNK | N/A | N/A |
|  | I13 | 9077 | 2185,016 | 194,391 | 1st | 2nd | Grandmother | 361,99 |
|  | I22 | 6645 | 1476,826 | 187,691 | 1st | 1st | Mother | 452,63 |

**Figure 2:**
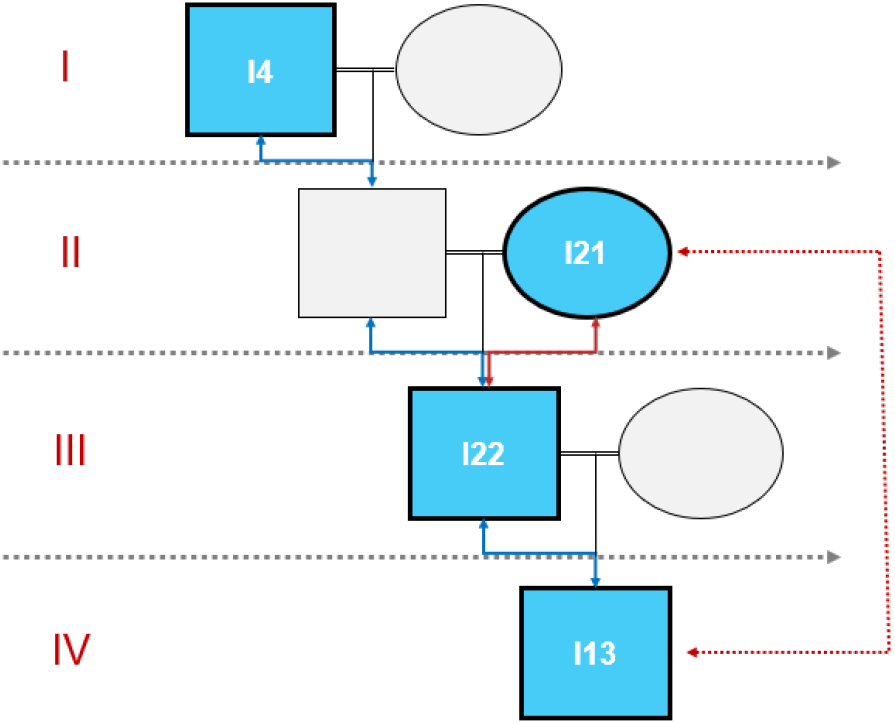
inferred familiypedigree. Family pedigree interred alter genetic study Roman numbers express the different generations. Squares represent males and circles females =: marriage ; 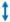 : patrilienal lineage with coincident Y-SNP markers; 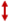 : shows relation with coincident X-SNP markers. 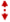 :shows relation with X-SNP markers inconsistencies.

## CONCLUSSION

Archaeological genetic studies have traditionally relied on uniparental markers to infer kinship within ancient burial contexts. Y-chromosome and mitochondrial DNA analyses have enabled the identification of shared paternal or maternal ancestry and have been successfully applied to historical cases such as Louis XVI [14]., the Romanov family [15]. and patrilineal lineages attributed to Genghis Khan [16]. However, while informative for assessing broad lineage continuity, these markers cannot resolve identifications or specific degrees of relatedness among individuals.

Autosomal STRs improve the ability to characterise close familial relationships and have been widely applied in forensic casework and mass-disaster victim identification [17]. Nonetheless, their resolving power declines sharply for more distant relationships and, crucially, their performance is severely compromised in highly degraded samples, as a result, historical or skeletal remains frequently yield partial profiles, allele dropout or stochastic artefacts that preclude reliable kinship inference [10,18,19]. These limitations have been repeatedly documented in archaeogenetic contexts and highlight the need for markers with greater tolerance to degradation. The development of autosomal SNP panels curated for kinship analyses has substantially increased resolution, and among these, the ForenSeq® Kintelligence kit represents an important advance. This targeted amplicon-sequencing approach interrogates 9,867 kinship-informative SNPs [13] and is optimised for low-template and degraded DNA, enabling reliable identity, ancestry and phenotype-related SNP calls from highly compromised samples.

Using this approach, we identified four members of the noble family of the Counts of Gelves and reconstructed a genealogical structure spanning four generations from the 17th–18th centuries (figure 2). The integration of autosomal SNP data with sex-chromosome markers reinforced pedigree assignments and allowed the discrimination of alternative kinship scenarios.

In one case, however, the UAS algorithm overestimated, the kinship degree, assigning a first-degree relationship between individuals I21 and I13, a result contradicted by X-chromosome evidence and the broader pedigree. This inflation is consistent with sister allele dropout in degraded samples, which reduces heterozygosity and increases opposing homozygous genotypes. To prevent zero likelihood ratios in these cases, UAS applies a corrective error-rate model that can inadvertently raise the combined LR when dropout is substantial. Beyond technical causes, the elevated consanguinity observed in this lineage likely contributed to reduced genome-wide heterozygosity—an expected consequence of consanguinity documented in both historical populations and noble lineages [20-22]. —and can similarly increase the probability of opposing homozygous genotypes, thereby affecting LR calculations. Consistent with this, the algorithm also inferred a third-degree relationship between individuals I4 and I21, a kinship supported by historical documentation and reflecting the progressive increase in consanguinity. A logLR was not computed for this pair, as additional historical sources are still being gathered to define the precise theoretical pedigree required for quantitative evaluation.

Despite the analytical challenges posed by degradation and consanguinity, our results show that Kintelligence can reliably recover autosomal SNP profiles from human remains nearly 500 years old and resolve extended familial relationships. This performance confirms the suitability of forensic SNP-based workflows for archaeological contexts and illustrates their value for reconstructing complex genealogical structures that cannot be addressed with traditional markers.

Most notably, this study demonstrates for the first time that an autosomal SNP approach routinely used in the forensic identification of living individuals can be successfully applied to archaeological material of early modern age. By analysing ∼10,000 kinship-informative SNPs, we reconstructed relationships up to the fourth degree and identified individuals with a level of confidence comparable to contemporary forensic casework. This achievement represents a qualitative leap for archaeological forensic genetics and establishes a robust foundation for future high-resolution kinship reconstruction in historical populations.

## ACKNOWLEGEMENTS

We thank the institutions that granted the necessary permissions to facilitate the archaeological, anthropological, and genetic work, as well as all team members whose collaboration made this study possible.

